## Supplementary material for "Improving the understanding of cytoneme-mediated morphogen gradients by *in silico* modeling"

### Supplementary Materials

#### Mathematical appendix

##### The dynamics of cytoneme lengths

$\lambda_r(t)$ and $\lambda_p(t)$ describe the dynamics of elongation and retraction of cytonemes emanating from either receiving $x_r$ or producing $x_p$ cell positions. It has been experimentally demonstrated that cytonemes have different dynamics (González-Méndez et al., 2017): they not only elongate and retract (Triangular behavior, Fig. 1E), but also have intermediate stationary phases, during which cytonemes maintain their maximum elongation (Trapezoidal behavior, Fig. 1F). Those behaviors have been implemented in this work as follows:

$$Triangular\ dynamics:$$

$$\lambda^{triangle}(t) = \begin{cases} v_e^{triang} \cdot t & if\ 0 \leq t \leq\ t_e^{triang} \\ v_{rt}^{triang}\left(t - t_e^{triang}\right) + \lambda_{max} & if\ \ t_e^{triang} < t \leq\ \ t_\tau^{triang} \end{cases} \quad \text{(Eq-SM1.1)}$$

$$Trapezoidal\ dynamics:$$

$$\lambda^{trapezoid}(t) = \begin{cases} v_e^{trap} \cdot t & if\ 0 \leq t <\ t_e^{trap} \\ \lambda_{max} & if\ t_e^{trap} \leq t <\ t_s + t_e^{trap} \\ v_{rt}^{trap}\left(t - (t_s + t_e^{trap})\right) +\ \lambda_{max} & if\ t_s + t_e^{trap} \leq t <\ t_\tau^{trap} \end{cases}$$

(Eq-SM1.2)

Where $v_e$ and $v_{rt}$ are the velocities of elongation and retraction for Triangular of Trapezoidal dynamics (described by superscripts *triang* and *trap* respectively). Both variables include the sign, being $sgn(v_e) = +1$ and $sgn(v_{rt}) = -1$.

$t_e$ is the time spent by a cytoneme in the elongation phase and $t_\tau$ is the total time from elongation to retraction; $t_\tau$ is defined as $t_\tau$ = $t_e$ + $t_{rt}$ for triangles and $t_\tau$ =$t_e$ + $t_s + t_{rt}$ for trapezoids, $t_s, t_{rt}$ being the time spent in the stationary and retraction phases respectively. $\lambda_{max}$ is the maximum length of a cytoneme (elongating with $v_e$ during $t_e$).

Expressions SM1 are defined for a time variable $t \in (t_0, t_\tau)$ with initial time $t_0 = 0$. In the code, cytonemes elongate and retract multiple times throughout the entire simulation; to this end, $t_{aux}$ (an auxiliary temporal variable) was introduced in the code for the SM1 equations.

The design of the temporal dependence of the contact functions (Eq-3) through the functions $\lambda(t)$ allows the introduction of different cytoneme dynamics, including the static case: $\begin{cases} \lambda_r(t) = \lambda_r \\ \lambda_p(t) = \lambda_p \end{cases}$

**The probability of contacts $\boldsymbol{\psi(\mu, x)}$**

The cellular mechanisms behind the establishment of cytoneme contacts are not well understood. We took advantage of the capability of Cytomorph to simulate situations difficult to study experimentally to predict the effect on the gradient of different types of contact functions $\psi(\mu, x)$. As an example, in what follows we describe the information obtained about this function looking at its dependence on different variables.

1. If $\psi(\mu, x)$ does not depend on cell position, then $\psi(\mu, x) = \psi(\mu)$. This implies that cytonemes always stablish contacts with a probability $\mu$, if they satisfy the distance condition needed to find another cell membrane (Eq-3). In this case, the probability of contact can be defined as the efficiency to establish connection for morphogen release.

$$\psi(\mu) = \begin{cases} 1\ with\ a\ probabilty\ of\ \mu \\ 0\ with\ a\ probabilty\ of\ 1-\mu \end{cases} \text{(Eq-SM2.1)}$$

Cytonemes overlapping for long stretches have experimentally been observed in *Drosophila* wing disc (González-Méndez et al., 2017). This can be modeled as a function independent of cell position assuming that new contacts along the overlapping stretch are more likely to occur after an initial contact between two cytonemes. For type 3 experimental case we developed a different module in the software in which the probability of function for $C_3(x_r)$ has been defined as follows:

$$\Psi(\mu) = \psi(\mu) + \delta_{\psi(\mu),1} \sum_{i=1}^{\lambda_{overlap}} \psi(\mu_i)$$

$$with\ \ \mu_i = \begin{cases} \mu + (r \cdot \mu) \cdot i & if\ \mu + (r \cdot \mu) \cdot i \ \leq\ 1 \\ 1 & if\ \mu + (r \cdot \mu) \cdot i \ >\ 1 \end{cases} \text{ and } \mu \in [0,1]$$

(Eq-SM2.2)

The contact probability for the overlapping $\mu_i$ increases with the number of previous contacts (i) in a rate $r$ that has to be determined experimentally (simulations in this work used $r = 0.2$).

Note that the single contact function is the same as in Eq-SM2.1, with the exception that the delta of Kronecker makes the sum different from zero only when a previous contact has been established. The sum depends on the integer overlapping length $\lambda_{overlap}$, which is a dynamic value determined by $\lambda_{overlap}(\mathrm{t}) = \lambda_\mathrm{r}(\mathrm{t}) + \lambda_\mathrm{p}(\mathrm{t}) - x_r - x_p$.

2. If $\psi(\mu, x)$ depends on the cell position within the tissue, this probability variable is related with the weight functions used in other models. These weight functions might indicate the existence of cellular mechanisms that control the establishment of a

contact with a specific cell to regulate the amount of transferred morphogen. This spatial dependence could also be related to the accessibility of the receiving cell, since cytonemes have to navigate through a tissue evading biochemical, mechanical and physical constraints.

In both possibilities the function can be written as:

$$\psi(\mu, x) = \psi(\mu(x)) = \begin{cases} 1 \; with\; a\; probabilty\; of\; \mu(x) \\ 0 \; with\; a\; probabilty\; of\; 1 - \mu(x) \end{cases} \text{(Eq-SM2.3)}$$

Where the function $\mu(x)$ could depend on the cellular mechanisms or on the tortuosity of the path. Since these possibilities are very specific of either the tissue or the path, in this work we implemented a simple case in which the probability decays linearly as $\mu = m \cdot x_r + n$.

The parameters m and n were calculated using the following conditions:

- The probability of contact $\mu$ is zero in regions more distant that the sum of the maximum elongation lengths of cytoneme protruding from the receiving and producing cells.

$$\mu(\lambda > \lambda_r + \lambda_p) = 0$$

- The probability function of contacts is normalized to 1.

$$\int_0^{\lambda_r+\lambda_p} \mu(x_r) dx_r = 1$$

Therefore, the final form used for the variable probability along the receiving cells was:

$$\mu(x_r) = \frac{2}{(\lambda_r+\lambda_p)} \left[ \frac{-x_r}{(\lambda_r+\lambda_p)} + 1 \right] \text{(Eq-SM.3)}$$

#### Cytomorph software architecture

Cytomorph was designed in different scripts that can be divided into the following modules (Supplementary Figure S.1):

1. A module containing a group of scripts for the graphical user interfaces (GUIs) was designed to run simulations.
2. A module to numerically simulate the cytoneme dynamics and to compute contacts and their spatial distribution over time.
3. A module to plot graphs to visualize the simulated spatial and temporal contact distribution and to understand the effects of different cytoneme features on gradient properties.

1. *Graphical user interface (GUI)*

The GUI is composed of 4 different scripts to create the graphical windows. ***Cytomorph*** is the main script and contains the code for the initial window (Fig. 2A-2) in which experimental data can be loaded and cytoneme features can be selected. If we want to compute different simulations and compare them, ***Cytomorph*** calls to the script ***cases2compare*** that creates a secondary window (Fig. 2B-1). In this window different combinations of parameters (cases) can be loaded to compare with the initial case, taken as reference. For convenience, if the cases to compare are just different values (scan) of the same variable, we also created a third script called ***Scan*** that generates a window (Fig. 2B-2) in which a scan of a variable value can be determined. Finally, there is another script called ***NewGraph*** that generates a window (Fig. 2B-3) in which the graphical properties can be selected if the user wants different options of default properties.

After collecting all the parameters, the main script ***Cytomorph*** calls the computing module to run the corresponding experimental data and the selected features to be studied.

2. *Numerical computation*

This module numerically simulates the cytoneme dynamics and computes the contacts and their distribution in the receiving cells for the parameters selected and for the experimental data loaded.

This module is made up of 6 different scripts, controlled by a main script (called ***CytomorphFunction***), which receives the parameters from the graphical module and calls the script functions according to the properties to be simulated.

If the case "dynamic cytonemes" is selected, then ***CytomorphFunction*** calls for the dynamic module. This module computes the sum of the possible contact distributions in the receiving cells, determined according to Eq-3 and Eq-SM1.

Since the mechanisms that influence cytoneme dynamics are not well understood, some hypotheses can be verified, such as the effect of the type of contact function $\psi(\mu, x)$. To this end we created a variable called CL; if selected (CL=1), the ***CytomorphFunction*** calls the script ***DynamicCL*** to compute the contact according to Eq-SM2.2. Otherwise (CL=0), the main script calls the script ***Dynamic*** that computes the contact distribution according to Eq-SM2.1, if the probability $\mu$ is a constant (case $\psi(\mu, x) = \psi(\mu)$), or to Eq-SM2.3, if the probability is a function of the position (case $\psi(\mu, x) = \psi(\mu(x))$).

Computationally $\psi(\mu)$ is coded by the function randsrc in Matlab as follows:

$$\psi(\mu) = randsrc(\,1, 1, [0\ \ 1; 1 - \mu\ \ \mu])$$

We observed that the computational runtime cost of this function of our code is high compared to other code sections. For convenience, if we are working with a contact probability of 100% ($\mu = 1$), to reduce the simulation time we created two more scripts called ***DynamicProb100*** and ***DynamicCLProb100*** in which we replaced function randsrc for $\psi(\mu) = 1$.

Finally, to also consider static cytonemes, we created a static module that computes the sum of all possible contact distributions in the receiving cells, determined according to Eq-3 for static cytonemes ($\lambda_{r,p}(t)$ =$\lambda_{r,p}$ =const). As in the dynamic case this module is divided in two different scripts: ***Static*** (CL=0) and ***StaticCL*** (CL=1) that compute the contact distribution according to the different SM2 equation.

*3. Plotting module*

While the main computation was done in the previous modules, all the information of the contact distribution was stored in an array to be subsequently used in the plotting module to depict different features that help the understanding of the software results.

This module is a main script called ***SimulationPlots*** in which the simulation results are graphically plotted in figures to facilitate the interpretation of the numerical simulations of our code. The selected magnitudes that Cytomorph plots as outputs are estimated as follows:

- *Contact distribution*

The software computes the number of contacts per cell along the simulated time. This is repeated over a wide range of simulations per case and the resulting contact distribution array is the base for computing the rest of the parameters. Since the software collects a random subgroup from the published experimental data values (González-Méndez et al., 2017) in each simulation, we can simulate the number of contacts of a specific receiving cell position ($x_r$) and see the predicted value per simulation (Fig. 2C-1). To visualize the contact distribution along simulations violin plots are shown in Fig. 2C-2.

- *Signal variability*

The predicted values for simulation variability can be determined from the different simulations. To better study this parameter and statistically compare different cases, we computed the distribution of coefficients of variation per case. To do this, we divided the N simulations runs into 100 subgroups of N/100 samples each. Then, the coefficient of variation distribution per case was performed over those 100 subgroups. The resulting data are presented as violin plots in Fig. 2C-3. (Note: if the number of simulation runs is lower than 1,000 (N<1,000), then the division is done into 10 subgroups of N/10 samples each).

- *Temporal evolution*

We can also observe the number of contacts in each receiving cell per time lapse (Fig. 2C-4) and the total evolution of the contact distribution and gradient shape over the simulated signaling time (Fig. 2C-5).

- *Gradient distribution*

The software computes the Eq-SM.5 in each iteration and subsequently, for a validation of the model, the software plots the simulation versus the experimental data of the morphogen gradient. In this way, we can visualize the accuracy of the *in silico* predictions.

As mentioned before, the software gives a numerical estimation for the number of contacts per receiving cell along time. The plotted gradient (Fig. 2C-6) is a normalized gradient calculated from Eq-SM.5; specifically, it is the normalized exponential fit of the numerical simulations, together with the standard deviation of the simulations performed per case (in error bars).

This plotting module also contains the next two auxiliary scripts:

***Violin:*** Since Violin plots are not implemented in the Matlab base code, we externalized these plots in the software using the script developed by Holger Hoffmann, which is available in MatlabWorks:

*Hoffmann H, 2015: violin.m - Simple violin plot using matlab default kernel density estimation. INRES (University of Bonn), Katzenburgweg 5, 53115 Germany.*

https://es.mathworks.com/matlabcentral/fileexchange/45134-violin-plot

***swtest***: Since Shapiro-Wilk test is not implemented in Matlab code, for the statistical study of normality we used the script developed by Ahmed Ben Saïda, which is available in MatlabWorks:

*Copyright (c) 17 March 2009 by Ahmed Ben Saïda,*

*Department of Finance, IHEC Sousse – Tunisia*

https://es.mathworks.com/matlabcentral/fileexchange/13964-shapiro-wilk-and-shapiro-francia-normality-tests

**Variability and fluctuations in Cytomorph.**

The inputs for the model via spreadsheet are the distributions of cytoneme lengths and elongation, stationary and retraction times for Triangle and Trapezoid cytoneme dynamics. Cytomorph randomly selects a subset of these data distributions for each simulation; this, together with the probability of contacts, generate small differences in the contact distribution for the simulated conditions, which is the source of variability and fluctuations in the model.

The variability of the model has been studied computing different parameters for each cell position (absolute number of contacts, relative number of contacts and coefficient of variation), developing different graphs to visualize the rough data (Supplementary Figure S.11 A-B) and plotting the summarized data in violin plots (Supplementary Figure S.11 C-D).

We were also concerned about the similarity between the predicted variability and the one experimentally observed. To compare with the measured experimental variability, in the final graphical representation of the gradient for each simulated condition we included the simulated variability (numerical standard deviation plotted in error bars)

In this work, we have also paid special attention to how this variability can be altered by different cytoneme features. Since we had observed that the relative variation in the numbers of contacts (standard deviation at position x divided by the mean number of contacts for that position) follows a general tendency to grow with receiving cell position (Supplementary Figure S.11 E), we selected the first cell row as the reference position to study the change in variability between different conditions. For that purpose, we have used graphs (Supplementary Figure S.11 A-D) and statistical tests (p-value matrices) as described in a previous section.

The largest fluctuations were found in the tail of the morphogen gradient (last rows of receiving cells). Since they correspond to the region in which the amount of morphogen is low, the fluctuations do not change the activation of the low-threshold targets like Cubitus interruptus (Ci). We then concluded that those fluctuations were biologically negligible for our study.

**Comparison of the normalized experimental data with the normalized contact function**

The concentration of morphogen released in each cell can be determined by the general equation:

$$[protein] = F(contact) - \delta \cdot [protein] \text{ (Eq-SM.4)}$$

where the function $F$ determines the quantity of morphogen transmitted in each contact; it depends on biological variables that control the morphogen transmission in the contact place. In this work, we assumed the all contacts are similar and therefore, each contact contributes with the same amount of morphogen. In this case, the function is the number of contacts per cell multiplied by a constant:

$$F(contact) = \alpha \cdot \#contact$$

As we want to compare the simulations with experimental data, we have first analyzed under which conditions it is possible to compare the following independent normalizations: the data of experimental fluorescent intensity (normalized to the maximum intensity $I_{max}$) versus the contact function in the theoretical model (normalized to the maximum contact function $\#contact_{max}$).

The equation SM.4 in a certain period of time $[t_i, t_{i+1}]$ can be written as:

$$[protein] = \alpha \cdot \#contact - \delta \cdot [protein] \cdot (t_{i+1} - t_i) \text{ (Eq-SM.5)}$$

If we normalize the equation SM.5 to the maximum concentration of morphogen we obtain:

$$\frac{[protein]}{[protein]_{max}} = \frac{\alpha \cdot \#contact}{[protein]_{max}} - \frac{\delta \cdot [protein] \cdot (t_{i+1} - t_i)}{[protein]_{max}}$$

$$\frac{[protein]}{[protein]_{max}}+\frac{\delta \cdot [protein]\cdot(t_{i+1}-t_i)}{[protein]_{max}}=\frac{\alpha \cdot \#contact}{[protein]_{max}}$$

$$(1+\delta \cdot (t_{i+1}-t_i))\cdot \frac{[protein]}{[protein]_{max}}=\frac{\alpha \cdot \#contact}{[protein]_{max}} \text{ (Eq-SM.6)}$$

Since it is a linear proportion ($\alpha$ = constant), the maximum intensity corresponds to the maximum number of contacts:

$$[protein]_{max}=\alpha \cdot \#conctact_{max}-\delta \cdot [protein]_{max}\cdot(t_{i+1}-t_i)$$

This can be approximated to $[protein]_{max} \approx \alpha \cdot \#conctact_{max}$ in the region of the maximum value, since we can consider $\alpha \cdot \#conctact_{max} \gg \delta \cdot [protein]_{max} \cdot (t_{i+1} - t_i)$.

Therefore, Eq-SM.6 can be written as:

$$\left(1+\delta \cdot (t_{i+1}-t_i)\right)\frac{[protein]}{[protein]_{max}} \approx \frac{\alpha \cdot \#contact}{\alpha \cdot \#conctact_{max}}=\frac{\#contact}{\#conctact_{max}} \text{ (Eq-SM.7)}$$

Since the fluorescence intensity of a GFP protein tagged to the morphogen (*Hh:GFP BAC*) was used to obtain the experimental data, the amount of protein is related to the intensity as $I=\beta \cdot [protein]$; then, if we normalize to the maximum intensity $I_{max}$ :

$$\frac{I}{I_{max}}=\frac{\beta \cdot [protein]}{\beta \cdot [protein]_{max}}=\frac{[protein]}{[protein]_{max}}$$

It is important to mention that the previous linear relation is only valid if the acquisition of image samples by confocal laser microscopy has been taken without reaching saturation and with a linear gamma function of acquisition.

Considering the previous confocal conditions, the normalization to the maximum contact in equation Eq-SM.7 can be compared with the experimental data (normalized to $I_{max}$) using the relation:

$$(1 + \delta \cdot (t_{i+1} - t_i))\frac{I}{I_{max}} \approx \frac{\#contact}{\#conctact_{max}} \text{ (Eq-SM.8)}$$

Therefore, the conditions to compare *in silico* simulations with experimental data are:

- Mathematically: The degradation rate of the morphogen should be taken into account using Eq-SM.8.
- Experimentally: Confocal images must have been taken according to the linear gamma function and within the limits of the acquisition range.

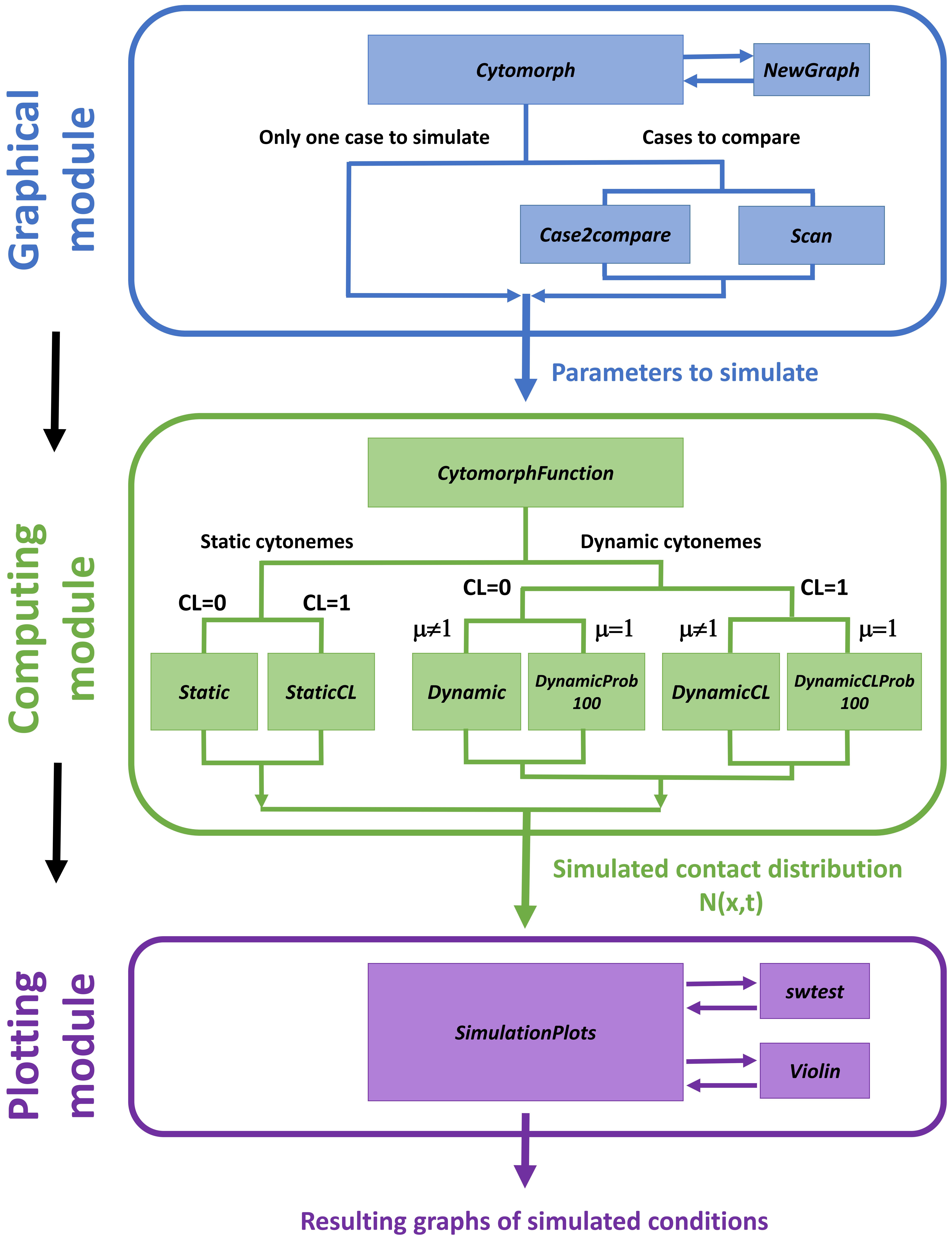
Graphical module
Cytomorph
NewGraph
Only one case to simulate
Cases to compare
Case2compare
Scan
Parameters to simulate
Computing module
CytomorphFunction
Static cytonemes
Dynamic cytonemes
CL=0
CL=1
CL=0
CL=1
μ≠1
μ=1
μ≠1
μ=1
Static
StaticCL
Dynamic
DynamicProb 100
DynamicCL
DynamicCLProb 100
Simulated contact distribution N(x,t)
Plotting module
SimulationPlots
swtest
Violin
Resulting graphs of simulated conditions

**Figure S.1 Schematic view of computational steps of Cytomorph machinery.** Cytomorph is composed by 14 different scripts that can be divided in three groups according to their function: In blue, scripts that code the graphical user interphase. In green, scripts that compute the dynamic of cytonemes and their contacts depending on different variables (static or dynamic cytonemes, if cytonemes can contact along their length (CL=1) or not (CL=0) and probability of contacts $\mu$). In purple, the scripts that plot the numerical simulations.

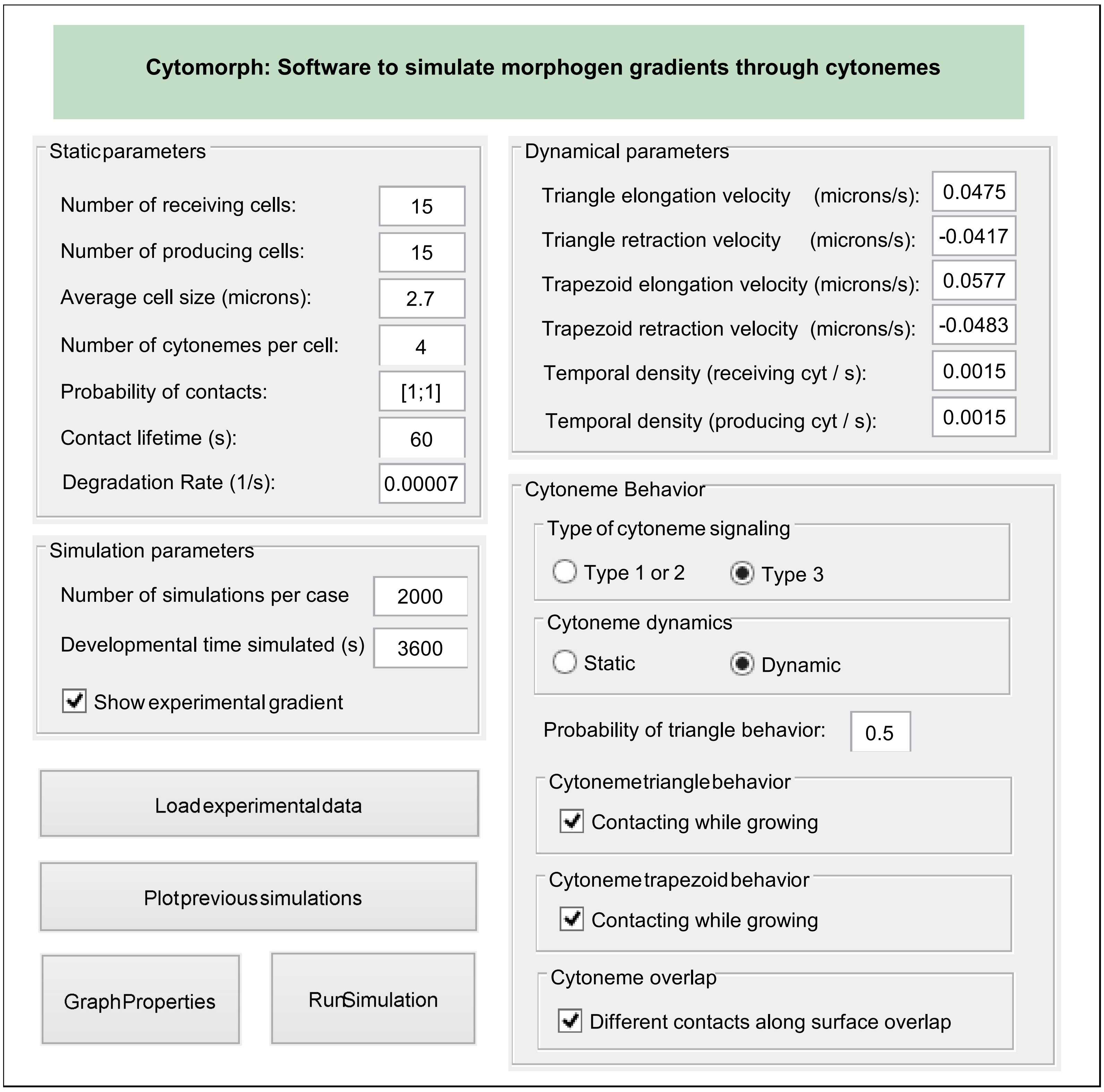
Cytomorph: Software to simulate morphogen gradients through cytonemes
Static parameters
Number of receiving cells:
15
Number of producing cells:
15
Average cell size (microns):
2.7
Number of cytonemes per cell:
4
Probability of contacts:
[1;1]
Contact lifetime (s):
60
Degradation Rate (1/s):
0.00007
Simulation parameters
Number of simulations per case
2000
Developmental time simulated (s)
3600
Show experimental gradient
Load experimental data
Plot previous simulations
Graph Properties
Run Simulation
Dynamical parameters
Triangle elongation velocity (microns/s):
0.0475
Triangle retraction velocity (microns/s):
-0.0417
Trapezoid elongation velocity (microns/s):
0.0577
Trapezoid retraction velocity (microns/s):
-0.0483
Temporal density (receiving cyt / s):
0.0015
Temporal density (producing cyt / s):
0.0015
Cytoneme Behavior
Type of cytoneme signaling
Type 1 or 2
Type 3
Cytoneme dynamics
Static
Dynamic
Probability of triangle behavior:
0.5
Cytoneme triangle behavior
Contacting while growing
Cytoneme trapezoid behavior
Contacting while growing
Cytoneme overlap
Different contacts along surface overlap

**Figure S.2. Main window of Cytomorph software.** Detailed view of the main GUI window of Cytomorph, in which different *in silico* conditions can be selected to study and simulate (with their respective units in brackets).

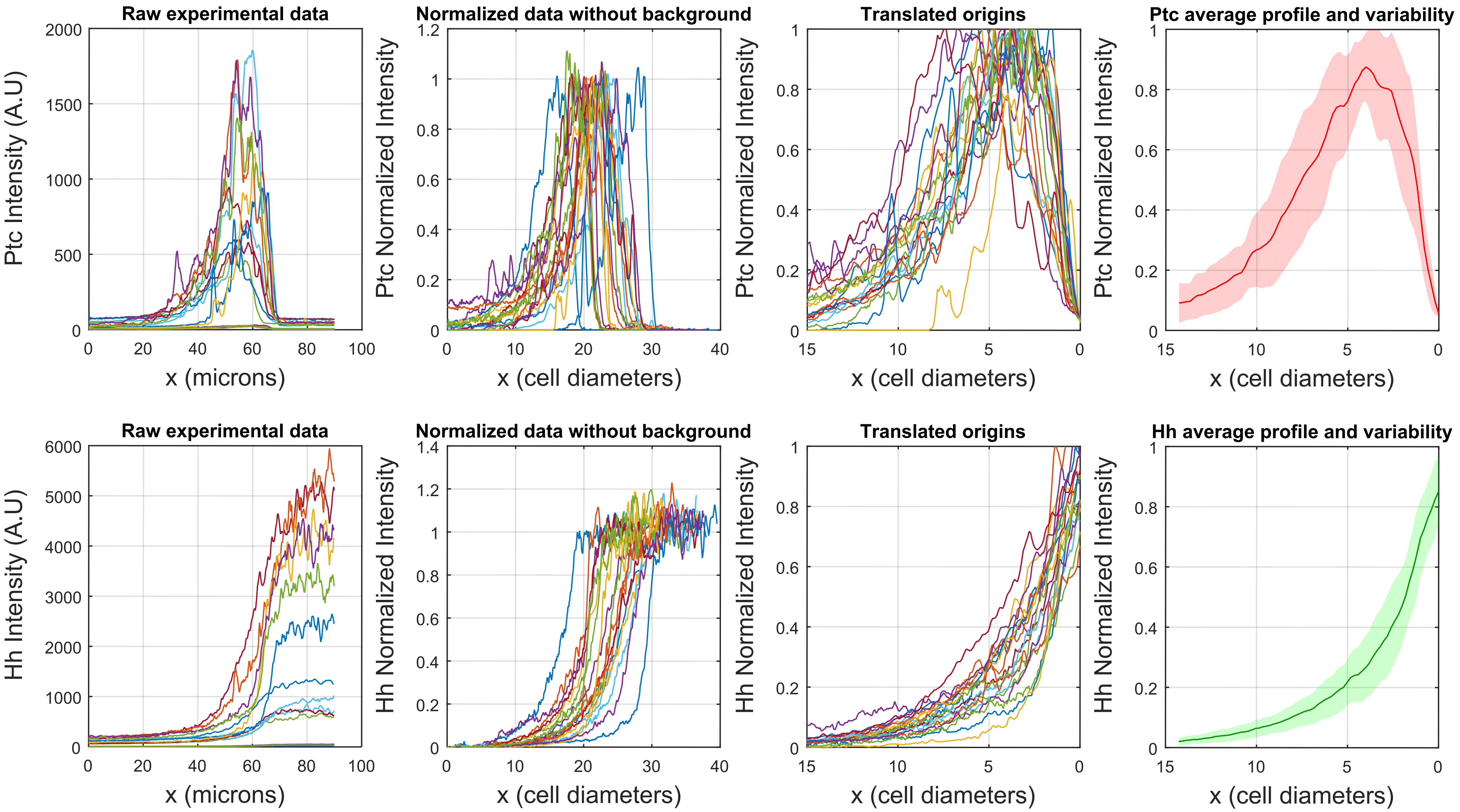
Raw experimental data
Ptc Intensity (A.U)
x (microns)
Normalized data without background
Ptc Normalized Intensity
x (cell diameters)
Translated origins
Ptc Normalized Intensity
x (cell diameters)
Ptc average profile and variability
Ptc Normalized Intensity
x (cell diameters)
Raw experimental data
Hh Intensity (A.U)
x (microns)
Normalized data without background
Hh Normalized Intensity
x (cell diameters)
Translated origins
Hh Normalized Intensity
x (cell diameters)
Hh average profile and variability
Hh Normalized Intensity
x (cell diameters)

**Figure S.3. Graphical representation of mathematical treatment for Hh and Ptc experimental expression profiles.** The normalization processes for Ptc (top) and Hh (bottom) profiles described in Materials and Methods are represented. The first column show the raw data obtained using the FIJI plot profile tool. The second column illustrates the normalized signal in cell diameters without background. The third column represents the same normalized data translated to the common origin at the A/P compartment border, using the beginning of Ptc expression to locate it. The final column shows the average (continuous thin lines) and the variability of the experimental samples (standard deviations are plotted in color shaded areas, Ptc in red and Hh in green).

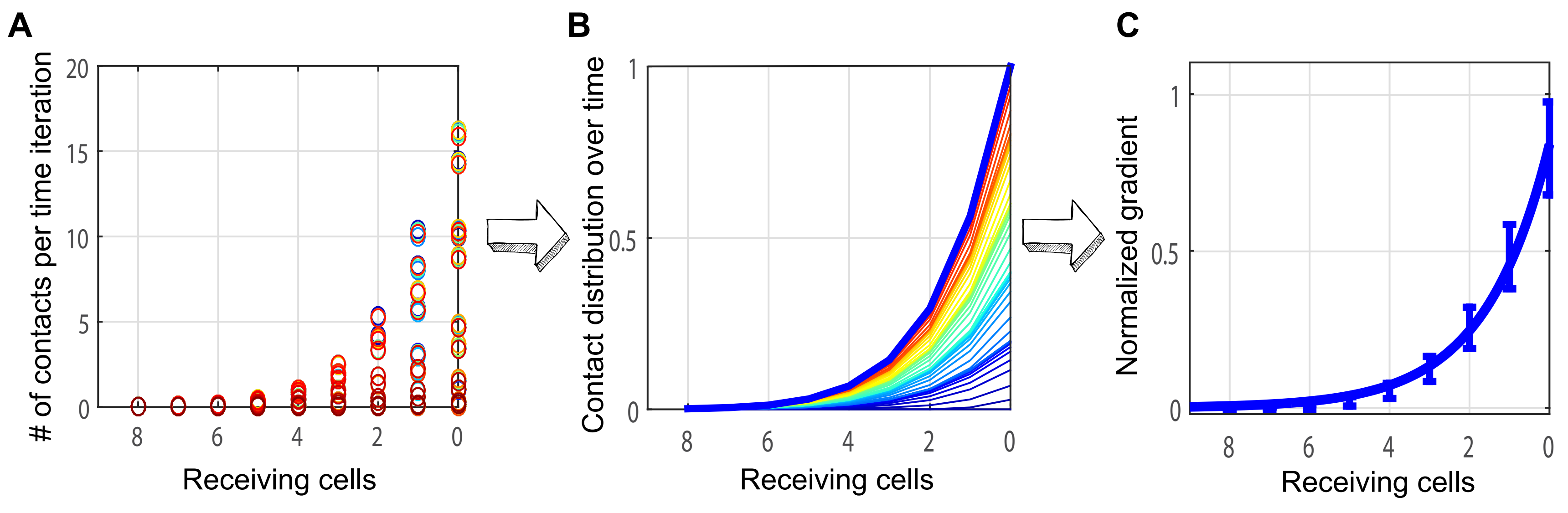
A
### of contacts per time iteration
20
15
10
5
0
8
6
4
2
0
Receiving cells
B
Contact distribution over time
1
0.5
0
8
6
4
2
0
Receiving cells
C
Normalized gradient
1
0.5
0
8
6
4
2
0
Receiving cells

**Figure S.4. The shape of the gradient is a consequence of the contact distributions.** The grafts show that the spatial distribution of contacts in the receiving cells (A) together with its temporal evolution (B) determine the final shape of the gradient (C).

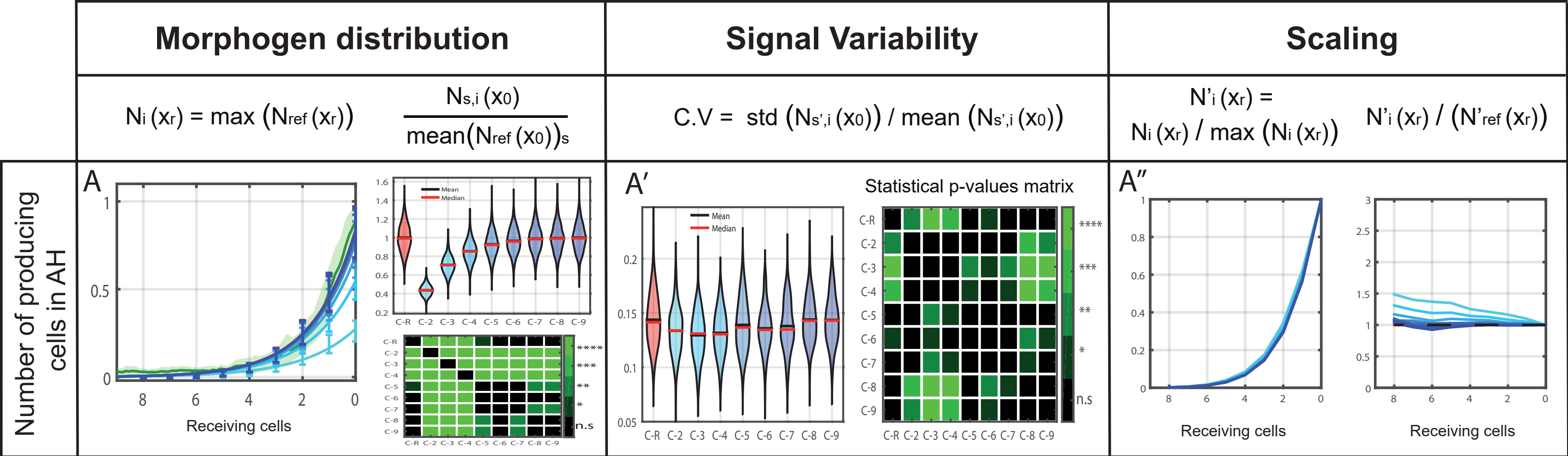

Morphogen distribution
Signal Variability
Scaling
Ni (xr) = max (Nref (xr))
Ns,i (x0) / mean(Nref (x0))s
C.V = std (Ns',i (x0)) / mean (Ns',i (x0))
N'i (xr) = Ni (xr) / max (Ni (xr))
N'i (xr) / (N'ref (xr))
Number of producing cells in AH
A
A′
A″
Receiving cells
Mean
Median
Statistical p-values matrix
C-R C-2 C-3 C-4 C-5 C-6 C-7 C-8 C-9
****
***
**
*
n.s

**Figure S.5. Influence of the number of producing cell rows in the shape of the gradient in abdominal histoblast nests.** The grafts show how different number of producing cell rows change the gradient properties: amount of the transmitted morphogen (A), signal variability (A') and scaling (A''). Since the length of the cytonemes and the size of the Hh producing cells in abdominal histoblast nets are smaller than those in imaginal wing discs, these results confirm the behavior observed in imaginal wing discs but with a scaling factor.

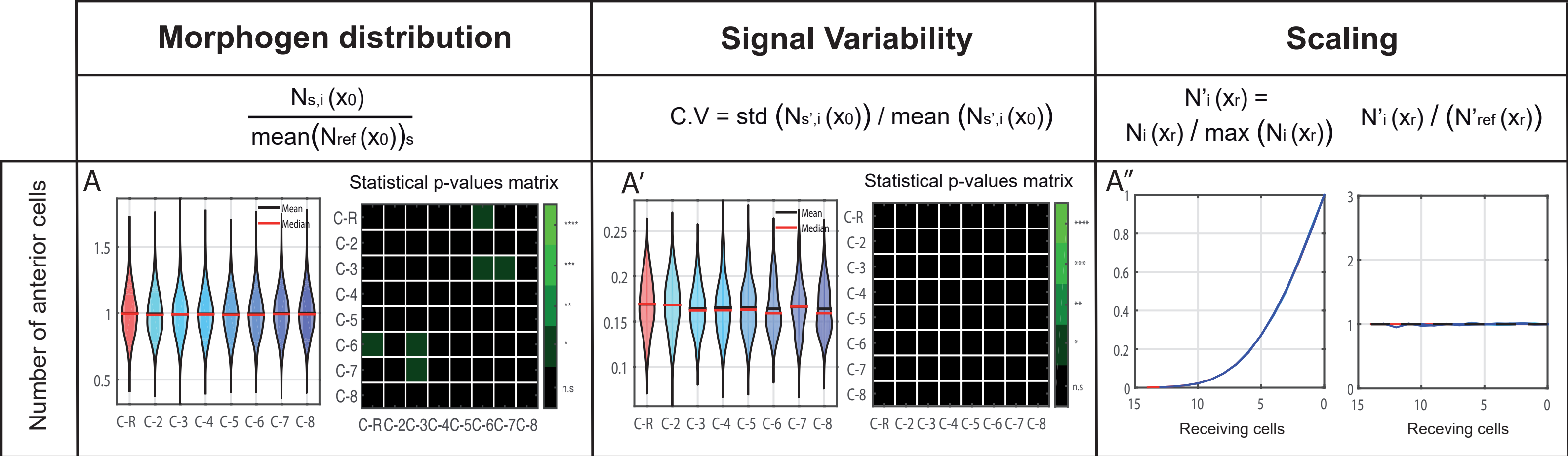

Morphogen distribution
Signal Variability
Scaling
Ns,i (x0) / mean(Nref (x0))s
C.V = std (Ns',i (x0)) / mean (Ns',i (x0))
N'i (xr) = Ni (xr) / max (Ni (xr))
N'i (xr) / (N'ref (xr))
Number of anterior cells
A
A′
A″
Statistical p-values matrix
Mean
Median
C-R
C-2
C-3
C-4
C-5
C-6
C-7
C-8
****
***
**
*
n.s
Receiving cells
Receving cells

**Figure S.6. Influence of the number of receiving cell rows in the shape of the gradient.** These results show that increasing the number of A compartment cell rows does not change the gradient properties: amount of transmitted morphogen (A), signal variability (A’) and scaling (A’’). These results are not surprising since the amount of Hh taken by each receiving cell of the A compartment only depends on the contacts of that cell with the producing cells of the P compartment.

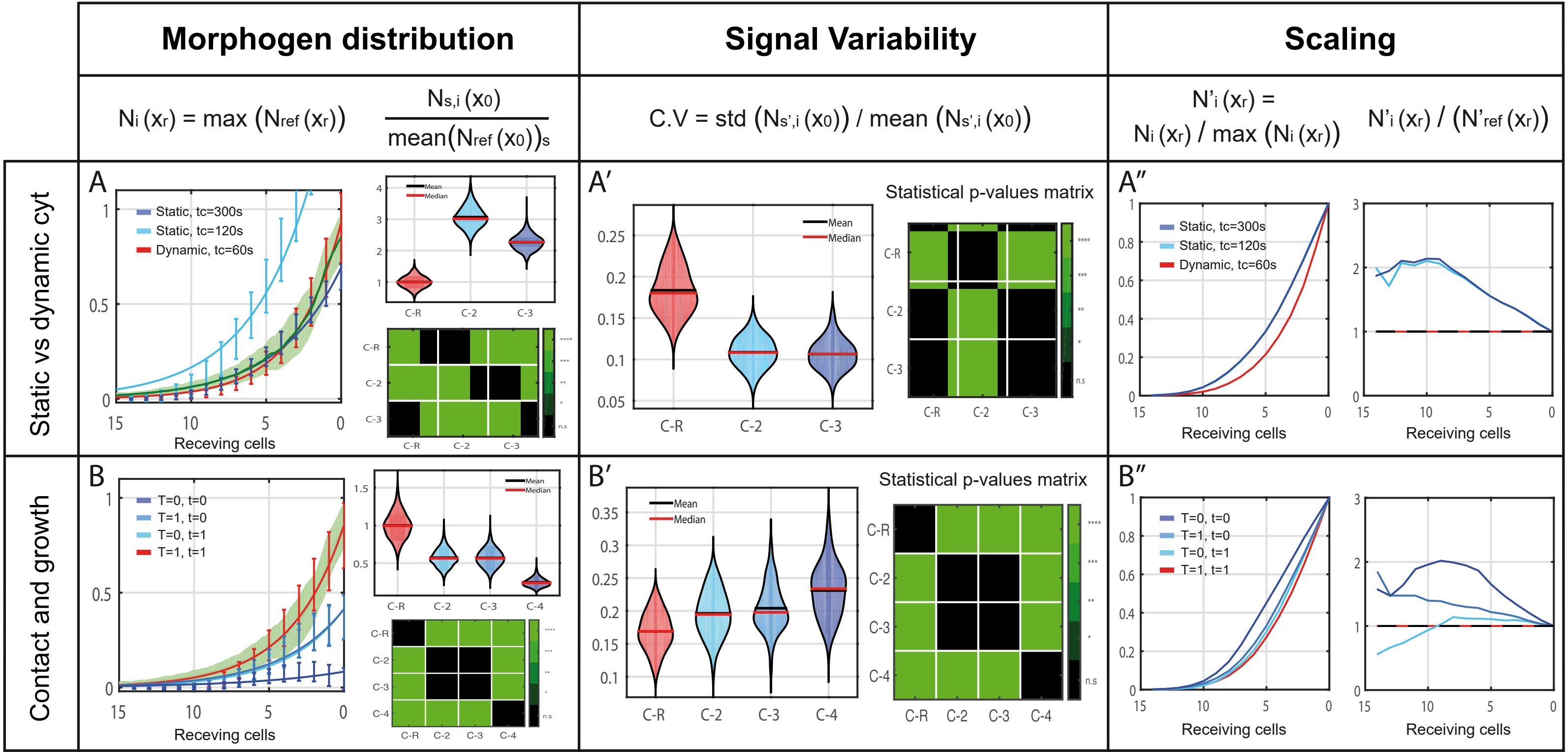

Morphogen distribution
Signal Variability
Scaling
Ni (xr) = max (Nref (xr))
Ns,i (x0) / mean(Nref (x0))s
C.V = std (Ns',i (x0)) / mean (Ns',i (x0))
N'i (xr) = Ni (xr) / max (Ni (xr))
N'i (xr) / (N'ref (xr))
Static vs dynamic cyt
Contact and growth
A
Static, tc=300s
Static, tc=120s
Dynamic, tc=60s
Receving cells
Mean
Median
C-R
C-2
C-3
A′
Statistical p-values matrix
A″
Receiving cells
B
T=0, t=0
T=1, t=0
T=0, t=1
T=1, t=1
C-4
B′
B″
****
***
**
*
n.s

**Figure S.7. Influence of the coordination of cytonemes growth and contact dynamics in gradient formation.** A) Simulations showed that dynamic and static cytonemes generate different gradient shapes. The differences suggest that cytoneme dynamics play a pivotal role in shaping the Hh gradient (A''). Simulations for static and dynamic cytonemes (dynamic with "tc=60s" in red, static with "tc=120s" in light blue, static with "tc=300s" in dark blue) showed that the final amount of morphogen released depends on factors, such as the time taken for a contact to be effective (time-length contacts "tc"). B) Growth simulations of triangular (t) and Trapezoidal (T) dynamics of cytonemes contacting while growing (=1) or contacting at maximum elongation (=0) (Ref.case T=t=1 in red, case 2: T=0,t=1 in light blue, case 3: T=1,t=0 in blue, case 4: T=0,t=0 in dark blue). The simulations shown for contact during growth using each type of behavior, triangular or trapezoidal, result in similar effects without statistically significant differences (C2 and C3 in B, B 'and B' '). However, if both types of cytoneme behaviors happen together, either contacting while growing or just contacting after growth (C-R and C4 respectively), there are relevant differences in the amount of morphogen transferred (B), the signal variability (B') and the gradient shape (B'').

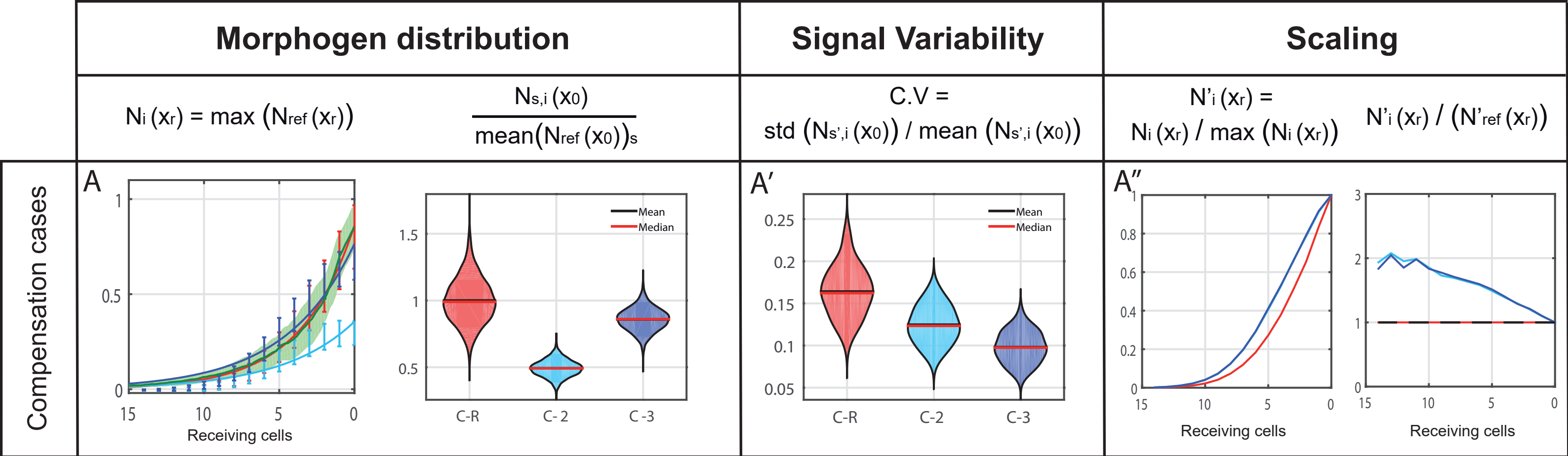
Morphogen distribution
Signal Variability
Scaling
Ni (xr) = max (Nref (xr))
Ns,i (x0) / mean(Nref (x0))s
C.V = std (Ns',i (x0)) / mean (Ns',i (x0))
N'i (xr) = Ni (xr) / max (Ni (xr))
N'i (xr) / (N'ref (xr))
Compensation cases
A
A'
A''
Receiving cells
Mean
Median
C-R
C-2
C-3

**Figure S.8. Signaling compensation between the number of producing cells and the number of cytonemes per cell.** Experimental data are represented in green, reference simulation for a normal tissue ($N_P$=15 and $n_{cyt}$=4) in red, altered tissue with less producing cells ($N_P$=3 and $n_{cyt}$=4) in light blue and altered tissue with a compensatory mechanism for this cell number reduction ($N_P$=3 and $n_{cyt}$=7) in dark blue. A) Left, morphogen distribution (y-axis) along the receiving cells normalized to the maximum value of the reference case. Right, violin plots for the number of contacts in the first row of receiving cell $x_0$, normalized to the reference average value (2000 simulations per case). A') Violin plots for the coefficient of variation (y-axis) per case (x-axis) in the first row of receiving cells ($x_0$). A'') Left, distribution of contacts normalized to their maximum (y-axis) to compare the changes in the shape in the receiving cells. Right, coefficient of the previous normalized distributions of contacts (y-axis) to study the scaling in the receiving cells.

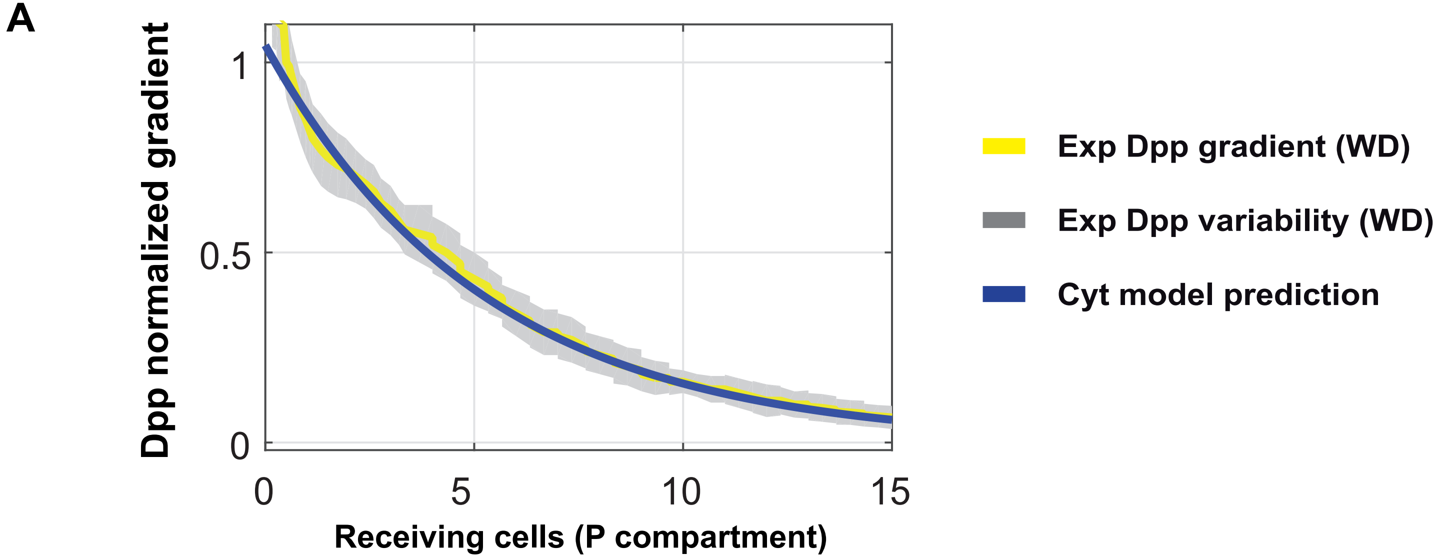


**B**

| | Morphogen distribution | Signal Variability | Scaling |
|---|---|---|---|
| | $N_i(x_r) = \max(N_{ref}(x_r))$ $\frac{N_{s,i}(x_0)}{\mathrm{mean}(N_{ref}(x_0))_s}$ | $C.V = \mathrm{std}(N_{s',i}(x_0)) / \mathrm{mean}(N_{s',i}(x_0))$ | $N'_i(x_r) = N_i(x_r) / \max(N_i(x_r))$ $N'_i(x_r) / (N'_{ref}(x_r))$ |
| Contact while growing | 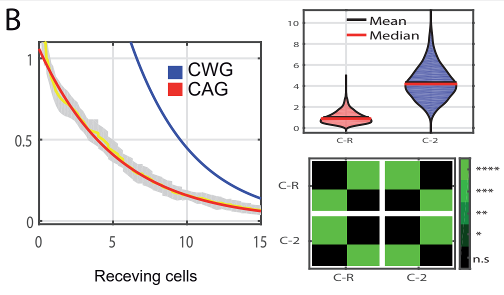 | 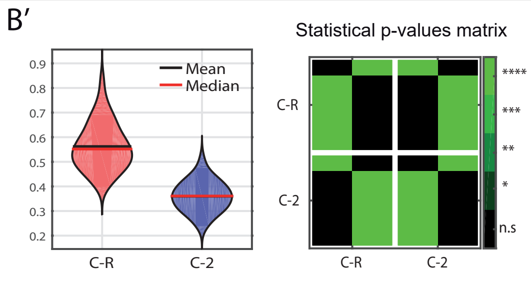 | 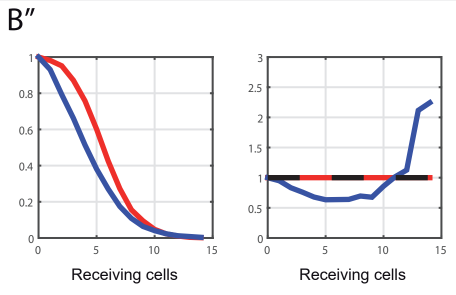 |

**Figure S.9. Cytomorph simulations for Dpp gradient in imaginal wing disc of Drosophila.** A) Cytomorph simulations for Dpp gradient (blue) compared to the real experimental Dpp gradient (Bollenbach et al., 2008)(average in yellow and experimental variability in grey). B) Dpp cytonemes contacting while growing (CWG, blue) or contacting after growth (CAG, red). The simulations show that two types of dynamics predict relevant differences in the amount of morphogen transferred (B), the signal variability (B') and the gradient shape (B''). Our results suggest that cytonemes contacting after growth fits better the experimental Dpp gradient.

Diffusion1D.m

```matlab
function [u]=Diffusion1D
%% Simulation Parameters

P(1) = 0.033; % Diffusion coefficient D
P(2) = 1; %c0
P(3)=0.00007; % Degradation rate
P(4)=1; % Eponential decay of the degradation term

L = 45; %Length of domain
tmax = 3600; %Max. simulation time
phi=3; %Cell diameter conversion factor (micras/cell)

m = 0; %Parameter corresponding to the symmetry of the problem (see help)
t = linspace(0,tmax,100); %tspan
x = linspace(0,L,100); %xmesh


%% solving with PDEPE

sol = pdepe(m,@DiffusionPDEfun,@DiffusionICfun,@DiffusionBCfun,x,t,[],P);
u = sol;
x=x/phi;

figure;
hold all

for n = linspace(1,length(t),10)
plot(x,u(n,:),'LineWidth',2)

end

xlabel('Distance in cell diameters','fontsize',18,'fontweight','b','fontname','arial')
ylabel('Normalized concentration','fontsize',18,'fontweight','b','fontname','arial')
axis([0 L/phi 0 P(2)])
set(gca,'XDir','Reverse','FontSize',15,'fontweight','b','fontname','arial')
grid on
```

DiffusionPDEfun.m

```matlab
function [c,f,s] = DiffusionPDEfun(x,t,u,dudx,P)
% Function defining the PDE
% Extract parameters
D = P(1);
deg=P(3);
n=P(4);
% PDE
c = 1;
f = D.*dudx;
s = -deg*(u.^n);
```

DiffusionBCfun.m

```matlab
function [pl,ql,pr,qr] = DiffusionBCfun(xl,ul,xr,ur,t,P)
% Boundary conditions for x = 0 and x = L;
% Extract parameters
c0 = P(2);
% BCs: No flux boundary at the right boundary and
% constant concentration on % the left boundary
pl = ul-c0; ql = 0; pr = 0; qr = 1;
```

DiffusionICfun.m

```matlab
function u0 = DiffusionICfun(x,P)
% Initial conditions for t = 0; can be a funciton of x
u0 = 0;
```

Diffusion1D Ln 18 Col 22

**Figure S.10. Matlab resolution of diffusion-degradation equation.** Screenshot of the computational code used for the simulation of the diffusion-degradation model. Left) Main script with *pdepe* function for 1-D parabolic and elliptic PDEs. Right) Three auxiliary functions called by *pdepe* that contain: the diffusion-degradation equation (top), the boundary conditions (middle) and the initial conditions (bottom) as described in Material and Methods.

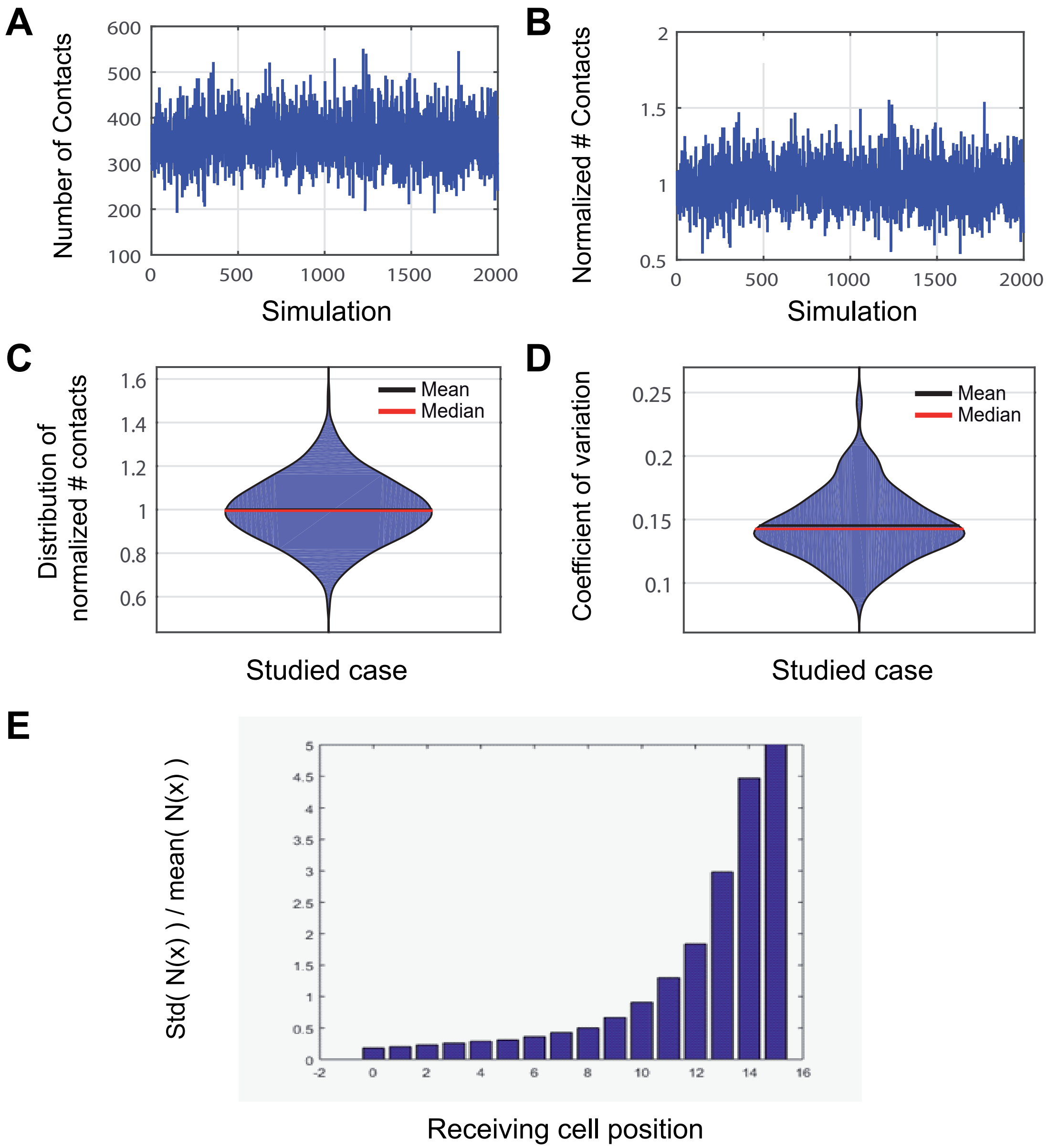

A
Number of Contacts
Simulation
B
Normalized # Contacts
Simulation
C
Distribution of
normalized # contacts
Mean
Median
Studied case
D
Coefficient of variation
Mean
Median
Studied case
E
Std( N(x) ) / mean( N(x) )
Receiving cell position

**Figure S.11. Study of the variability and fluctuations in the model.** The figure shows different representations of the variability simulated by Cytomorph: A) Total number of contacts per simulation in the first cell row, $x_0$. B) Number of contacts per simulation in the first cell row ($x_0$) normalized to the mean value over all simulations. C) Violin plot of data distribution in the image B. D) Violin plot of the coefficients of variation of the previous data. E) Simulated relative variability per cell position: $(std(N(x))/mean(N(x)))$.

#### Supplementary movies

**Movie 1. Cell organization at the A/P compartment border in abdominal histoblast nests.** It shows a confocal Z-stack from apical to basal of an A2 abdominal histoblast nest of a *Drosophila* pupa, with the A compartment marked with life-actin-RFP (Red) and the P compartment marked with CD8GFP (Green). Scale bar: 10 µm.

**Movie 2. *In vivo* movie of wild type abdominal histoblast cytonemes.** *In vivo* abdominal histoblast cytonemes located at the basal side of the tissue; confocal sections were taken at one-minute interval. The left movie shows merge channels of A compartment cells labeled with life-actin-RFP (in red) and of P compartment cells are marked with CD8GFP (in green). The movie located in the middle shows a single channel corresponding to A compartment cytonemes, and the right movie shows a single channel corresponding to P compartment cytonemes. Scale bar: 15 µm.

**Movie 3. Abdominal histoblast nest proliferation and migration.** While migration and proliferation of dorsal A and P compartment histoblast nests take place, larval epidermal cells are eliminated. The protein marked *Hh:GFP BAC* allows the visualization of Hh in the producing cells (P compartment histoblast nest, in green). The *EnhancerPtcRed* reporter correspond to *ptc* transcriptional activity in response of Hh gradient; this allows the visualization of the Hh gradient response in the receiving cells (A compartment histoblast nests, in red). Scale bar: 30 µm.

#### Supplementary tables

| X gradient (cell diameters) | Normalized Gradient signal | Signal variability ( Exp std ) | Receiving cytoneme length (μm) | Producing cytoneme length (μm) | Receiving elongation times for triangles (s) | Receiving retraction times for triangles (s) | Receiving elongation times for trapezoids (s) | Receiving stationary times for trapezoids (s) | Receiving retraction times for trapezoids (s) | Producing elongation times for triangles (s) | Producing retraction times for triangles (s) | Producing elongation times for trapezoids (s) | Producing stationary times for trapezoids (s) | Producing retraction times for trapezoids (s) |
|---|---|---|---|---|---|---|---|---|---|---|---|---|---|---|
| Value 1 | Value 1 | Value 1 | Value 1 | Value 1 | Value 1 | Value 1 | Value 1 | Value 1 | Value 1 | Value 1 | Value 1 | Value 1 | Value 1 | Value 1 |
| Value 2 | Value 2 | Value 2 | Value 2 | Value 2 | Value 2 | Value 2 | Value 2 | Value 2 | Value 2 | Value 2 | Value 2 | Value 2 | Value 2 | Value 2 |
| … | … | … | … | … | … | … | … | … | … | … | … | … | … | … |
| Value N | Value N | Value N | Value N | Value N | Value N | Value N | Value N | Value N | Value N | Value N | Value N | Value N | Value N | Value N |

**Table S.T 1 Units of the model.** Distribution of parameters (with their respectively units) that should be updated by an Excel file to Cytomorph.

#### Table S.2 Parameters used in cytoneme model simulations

| Simulations and cases | $N_R$ | $N_P$ | Average Cell size Φ (μm) | Number of simulations | Probability of contact μ | Signaling type<br>Type1-2=0<br>Type3=1 | Contact along overlap surface on? | Ncyt | Degradation rate ($s^{-1}$) | Time of development to simulate (s) | Time of contact $t_c$(s) | Dynamic on? | ve triangle (cell/s) | vrt triangle (cell/s) | ve Trapezoid (cell/s) | vrt Trapezoid (cell/s) | Probability of having triangles $p_t$ | Contact along growing on? | Exp data |
|---|---|---|---|---|---|---|---|---|---|---|---|---|---|---|---|---|---|---|---|
| Exp Vs Model WD | 15 | 15 | 3 | 2000 | [1;1] | 1 | 1 | 4 | $7\cdot10^{-5}$ | 3600 | 60 | 1 | 0.016 | -0.014 | 0.019 | -0.016 | 0.5 | T=t=1 | WD |
| Exp Vs Model AH | 9 | 9 | 4.37 | 2000 | [1;0.5] | 1 | 0 | 4 | $7\cdot10^{-5}$ | 3600 | 60 | 1 | 0.011 | -0.010 | 0.013 | -0.011 | 0.5 | T=t=1 | AH |
| Reference case | 15 | 15 | 3 | 2000 | [1;1] | 1 | 1 | 4 | $7\cdot10^{-5}$ | 3600 | 60 | 1 | 0.016 | -0.014 | 0.019 | -0.016 | 0.5 | T=t=1 | WD |
| Change in cell size | 15 | 15 | 2.5 to 3.5 each 0.2 | 2000 | [1;1] | 1 | 1 | 4 | $7\cdot10^{-5}$ | 3600 | 60 | 1 | 0.048/Φ | -0.042/Φ | 0.057/Φ | -0.048/Φ | 0.5 | T=t=1 | WD |
| Density of cytonemes | 15 | 15 | 3 | 2000 | [1;1] | 1 | 1 | 1 to 3 each 1 | $7\cdot10^{-5}$ | 3600 | 60 | 1 | 0.016 | -0.014 | 0.019 | -0.016 | 0.5 | T=t=1 | WD |
| Number of producing cells | 15 | 1 to 14 each 1 | 3 | 2000 | [1;1] | 1 | 1 | 4 | $7\cdot10^{-5}$ | 3600 | 60 | 1 | 0.016 | -0.014 | 0.019 | -0.016 | 0.5 | T=t=1 | WD |
| Number of receiving cells | 2 to 14 each 2 | 15 | 3 | 2000 | [1;1] | 1 | 1 | 4 | $7\cdot10^{-5}$ | 3600 | 60 | 1 | 0.016 | -0.014 | 0.019 | -0.016 | 0.5 | T=t=1 | WD |
| Compensation cases | 15 | 3 | 3 | 2000 | [1;1] | 1 | 1 | C2=4<br>C3=7 | $7\cdot10^{-5}$ | 3600 | 60 | 1 | 0.016 | -0.014 | 0.019 | -0.016 | 0.5 | T=t=1 | WD |
| Type of signaling: Type 3 | 15 | 15 | 3 | 2000 | [1;1] | 1 | 1 | 4 | $7\cdot10^{-5}$ | 1800 | 60 | 1 | 0.016 | -0.014 | 0.019 | -0.016 | 0.5 | T=t=1 | WD |
| Type of signaling: Type 1-2 | 15 | 15 | 3 | 2000 | [1;1] | 0 | 1 | 4 | $7\cdot10^{-5}$ | 1800 | 60 | 1 | 0.016 | -0.014 | 0.019 | -0.016 | 0.5 | T=t=1 | WD |
| No overlapping cytonemes | 15 | 15 | 3 | 2000 | [1;1] | 1 | 0 | 4 | $7\cdot10^{-5}$ | 3600 | 60 | 1 | 0.016 | -0.014 | 0.019 | -0.016 | 0.5 | T=t=1 | WD |
| Probability function | 15 | 15 | 3 | 2000 | Linear decay | 1 | 0 | 4 | $7\cdot10^{-5}$ | 3600 | 60 | 1 | 0.016 | -0.014 | 0.019 | -0.016 | 0.5 | T=t=1 | WD |
| Probability of contact | 15 | 15 | 3 | 2000 | [0.4;0.4] to [0.8;0.8] each [0.2;0.2] | 1 | 1 | 4 | $7\cdot10^{-5}$ | 3600 | 60 | 1 | 0.016 | -0.014 | 0.019 | -0.016 | 0.5 | T=t=1 | WD |
| Overlapping cytonemes | 15 | 15 | 3 | 2000 | [1;1] | 1 | 1 | 4 | $7\cdot10^{-5}$ | 3600 | 60 | 1 | 0.016 | -0.014 | 0.019 | -0.016 | 0.5 | T=t=1 | WD |
| Contact and growth | 15 | 15 | 3 | 2000 | [1;1] | 1 | 1 | 4 | $7\cdot10^{-5}$ | 3600 | 60 | 1 | 0.016 | -0.014 | 0.019 | -0.016 | 0.5 | C2:T=0,t=1<br>C3:T=1,t=0<br>C4:T=t=0 | WD |
| FRAP simulation | 7 | 7 | 4.37 | 2000 | [1;0] | 0 | 0 | 4 | $7\cdot10^{-5}$ | 3360 | 60 | 1 | 0.011 | -0.010 | 0.013 | -0.011 | 0.5 | T=t=1 | AH |
| Cytoneme triang/trap dynamics | 15 | 15 | 3 | 2000 | [1;1] | 1 | 1 | 4 | $7\cdot10^{-5}$ | 3600 | 60 | 1 | 0.016 | -0.014 | 0.019 | -0.016 | C2=0.1<br>C3=0.9 | T=t=1 | WD |
| Static vs dynamic: Dynamical case | 15 | 15 | 3 | 2000 | [1;1] | 1 | 0 | 4 | $7\cdot10^{-5}$ | 1800 | 60 | 1 | 0.016 | -0.014 | 0.019 | -0.016 | 0.5 | T=t=1 | WD |
| Static vs dynamic: static | 15 | 15 | 3 | 2000 | [1;1] | 1 | 0 | 4 | $7\cdot10^{-5}$ | 1800 | C1=120<br>C2=300 | 0 | 0.016 | -0.014 | 0.019 | -0.016 | 0.5 | T=t=1 | WD |
| Number of producing cells AH | 9 | 1 to 8 each 1 | 4.37 | 2000 | [1;0.5] | 1 | 0 | 4 | $7\cdot10^{-5}$ | 3600 | 60 | 1 | 0.011 | -0.010 | 0.013 | -0.011 | 0.5 | T=t=1 | AH |
| Dpp exp Model WD | 15 | 7 | 3 | 2000 | [1;1] | 1 | 1 | 4 | $2.52\cdot10^{-4}$ | 600 | 60 | 1 | 0.019 | -0.016 | 0.016 | -0.014 | 0.5 | T=t=0 | WD |

**Table S.2 Parameters used in cytoneme model simulations.** Numerical parameters used in the cytoneme model for each simulation. When a scan of a single variable is performed, the values are written from the initial to the final values with a specific step between them. Binary values mean that this condition is on=1 or off=0. WD and AH are abbreviations for wing imaginal discs and abdominal histoblast nests and T and t for Trapezoidal and Triangular behaviors respectively. Units are specified at the top of each column**.**

| k-s test | | Anterior | | | | |
|---|---|---|---|---|---|---|
| | | ts | te | | tr | |
| Posterior | ts | ns | Triangle | Trapezoid | Triangle | Trapezoid |
| | te | Triangle | ns | 0.027 | ns | 0.02 |
| | | Trapezoid | 0.015 | ns | 0.001 | ns |
| | tr | Triangle | ns | 0.0014 | ns | 0.011 |
| | | Trapezoid | ns | ns | 0.064 | ns |

| k-s test | | Anterior | | | | |
|---|---|---|---|---|---|---|
| | | ts | te | | tr | |
| Anterior | ts | | Triangle | Trapezoid | Triangle | Trapezoid |
| | te | Triangle | | 0.035 | ns | 0.044 |
| | | Trapezoid | | | 0.0028 | ns |
| | tr | Triangle | | | | 0.02 |
| | | Trapezoid | | | | |

| k-s test | | Posterior | | | | |
|---|---|---|---|---|---|---|
| | | ts | te | | tr | |
| Posterior | ts | | Triangle | Trapezoid | Triangle | Trapezoid |
| | te | Triangle | | 0.022 | ns | ns |
| | | Trapezoid | | | 0.0005 | ns |
| | tr | Triangle | | | | 0.04 |
| | | Trapezoid | | | | |

**Table S.3 Statistical study of the phase times used to compare cytoneme dynamics.**

Statistical p-values used to compare elongation, retraction and stationary phases using a Kolmogorov-Smirnov statistical analysis (n.s = no significance).

| Tissue | Diffusion constant: D(μm$^2$/ s$^{-1}$) | Length of simulated domain (μm) | Cell diameter Φ (μm) | Developmental time (s) | Degradation rate: δ (s$^{-1}$) | Average experimental maximum (Source normalization): $\langle u_{exp}^{N} \rangle_{x=0}$ |
|---|---|---|---|---|---|---|
| **Wing disc** | 0.033 | 48 | 3 | 3600 | $7 \cdot 10^{-5}$ | 0.852 |
| **Abdominal histoblasts** | 0.033 | 39.33 | 4.37 | 3600 | $7 \cdot 10^{-5}$ | 0.878 |
| **Abdominal histoblasts** | 0.011 | 39.33 | 4.37 | 3600 | $7 \cdot 10^{-5}$ | 0.878 |

**Table S.4 Diffusion parameters used in each simulation.** Numerical parameters used in the Figure 4 for diffusion simulations. Units are specified at the top of each column.
